# Striatal Dopamine Modulates Temporal Surprise P3a

**DOI:** 10.1101/2024.03.05.583515

**Authors:** Grace A. Whitaker, Michael Schwartze, Sarah Martin, Aland Astudillo, Monty Silverdale, Wael El-Deredy, Sonja A. Kotz

## Abstract

Dopamine is vital in forming mental models of “what” and “when” sensory events occur that essentially guide goal-directed behaviour. However, it remains largely unknown how variations in temporal predictability are incorporated into such mental models. A better understanding of the underlying mechanisms is important, considering dopaminergic depletion in diseases such as Parkinson’s Disease and Schizophrenia, where abnormal temporal processing is observed. Some electroencephalographic (EEG) studies indicate that noradrenergic mechanisms, as reflected in the P3b event-related potential, are modulated by temporal predictability, whereas others indicate that dopaminergic mechanisms as reflected in the P3a, underlie surprise. In this study, resting-state and task-dependent EEG was recorded from 24 healthy participants who were administered a selective D2 agonist or antagonist before they performed a pure tone auditory “oddball” task. Two oddball sequences included either partially predictable, with inter-stimulus intervals (ISIs) of 400ms/1200ms; or fully temporally predictable tones, with a consistent ISI of 600ms. Tones following 400ms ISIs were perceived as surprising, or “early”, as shown in an enhanced P3a response; tones following a 1200ms ISIs showed a much reduced P3a response (“late”). The agonist accentuated the “late” effect, demonstrating that drugs targeting D2 receptors modulate temporal prediction. These findings differentiate the role of the dopaminergic system in temporal processing and model-based auditory predictions.

## 1 Introduction

Being able to predict “what” and “when” events occur in the environment is an important ability to guide efficient goal-directed cognitive behaviour. For instance, individuals predict their turn to speak in a conversation based on the length of the final syllables before a pause, the length of the pause, and changes in pitch (Ford & Thompson, 1996; Wennerstrom & Siegel, 2003; Zellner, 1994). Similarly, musicians synchronise their performances based on prior knowledge of the length of pauses and the rhythmic structure of music (Fonseca-Mora et al., 2011; Koelsch, 2011). When the expected stimulus timing deviates in these situations, it may increase uncertainty in perception, leading to problems with coordination (Ford & Thompson, 1996; Large, 2008). Accordingly, humans must factor time into their mental models of the environment to efficiently predict events and to deal with varying degrees of temporal uncertainty (Buhusi & Meck, 2005; Ivry & Schlerf, 2008).

The updating of mental models is frequently studied using the oddball paradigm (Linden, 2005; Polich & Criado, 2006): Classical oddball experiments present a stream of repetitive standard stimuli (e.g., an equitone sequence) that is occasionally interrupted by deviant stimuli (e.g., change in frequency) (Squires, Squires, & Hillyard, 1975). Following the deviant stimulus, or “oddball event”, a prominent positive deflection of the event-related potential (ERP) of the electroencephalogram (EEG) is observed. This component reflects the processing of the stimulus, and peaks at around 300ms post-stimulus onset, and is termed the P3 (Donchin, 1981; Linden, 2005).

Using the oddball paradigm, studies have shown that deviance processing involves the registration of discrepancy between the expected and actual input and the updating of the mental representation of a stimulus (Polich, 2003; Polich & Criado, 2006). These different cognitive processes are associated with two subcomponents of the P3 complex; i.e. the P3a and P3b, respectively (Desimone et al., 1995; Knight, 1996; Squire & Kantrel, 1999). Their functional and topographical distinctions make the P3a and P3b useful tools for the delineation of attentional engagement and memory updating processes, respectively (Donchin, 1981; Polich & Criado, 2006).

The neurotransmitter dopamine is thought to play a crucial role in deviance processing, especially in the striatum of the basal ganglia. For example, the oddball P3a is reported to be modulated by dopaminergic genes (Heitland et al., 2013; Marco-Pallarés et al., 2010). Administration of the dopamine-2 (D2) receptor antagonist haloperidol has been shown to reduce the P3a amplitude (Kähkönen et al., 2002). Parkinson’s disease (PD) is characterised by the loss of dopamine-generating neurons in the substantia nigra, which supply the dorsal striatum with dopamine (Ayano, 2016). In PD, a reduced P3a amplitude has been proposed as a marker of disease progression (Seer et al., 2016; Solís-Vivanco et al., 2015); and dopaminergic treatments of PD can restore P3a modulation (Georgiev et al., 2015). Persons with Schizophrenia have abnormally high levels of striatal dopamine signalling, as evidenced by a reduced oddball P3a and P3b response compared to controls (Bramon et al., 2004; Yeon & Polich, 2003; Ford 1999).

These findings indicate that typical deviance processing requires balanced levels of striatal dopamine, as observed for other forms of cognition such as working memory, reversal learning, attention switching, and goal maintenance (Cools & D’Esposito, 2011; Cools et al., 2009; Wallace et al., 2011; Cools et al., 2007). Furthermore, the findings in Schizophrenia indicate that dopamine may influence both the initial registration of deviance (P3a), and the updating of a mental representation (P3b). This is somewhat at odds with previous studies showing that the P3b is more closely linked to noradrenaline than dopamine signalling (Nieuwenhuis, Aston-Jones, & Cohen, 2005).

One important study constraint is the reliance on patient populations, making it difficult to control for the heterogeneity in disorder severity, medication status, and the presence of comorbidities (Mushquash, Fawcett, & Klein, 2012). For example, the impairments observed in PD and Schizophrenia are not constrained to striatal dopamine signalling (Ayano, 2016; Gouzoulis-Mayfrank et al., 2007). As such, it is also difficult to draw conclusions how striatal dopamine affects deviance processing using patient studies alone.

Another important restriction of patient-based dopamine studies is that they are primarily concerned with deviance in “what” occurs in the environment, although efficient mental models must also be flexible to different degrees of temporal uncertainty (i.e., “when” deviance occurs). The importance of studying the role of dopamine regarding temporal uncertainty is highlighted by disorders such PD and Schizophrenia, where people with these disorders suffer debilitating impairments in auditory-motor synchronisation and temporal processing (Benoit et al., 2014; Dalla Bella et al., 2015; Woerd et al., 2018).

Temporal uncertainty can be studied by altering the timing of the stimulus presentation in an oddball paradigm. For example, it has been shown that the P3b amplitude increases when stimuli are temporally predictable rather than unpredictable (Schmidt-Kassow, Schubotz, & Kotz, 2009; Schwartze et al., 2011). In other words, deviance processing is enhanced when stimuli are presented regularly in time. This is thought to reflect the preferential processing of deviant stimuli at expected versus unexpected time points (Kotz, Schwartze, & Schmidt-Kassow, 2009; Schwartze et al., 2011). Dysfunctions in such preferential processing have been observed in patients with basal ganglia (Schwartze et al., 2011) and cerebellar lesions (Kotz, Stockert, & Schwartze, 2014), and are thought to affect the adaptation to a dynamic environment (Nozaradan et al., 2017; Schwartze, Keller, & Kotz, 2016). However, to date there has been no investigation on how dopamine alters deviance processing under varying degrees of temporal uncertainty.

In the present oddball study, dopamine was manipulated in healthy participants to investigate how dopamine over- and under-signalling influences temporal processing. The highly selective D2 receptor agonist cabergoline (Gerlach et al., 2003) and antagonist amisulpride (Tardieu et al., 2003) were administered, as D2 receptors are most abundant in the striatum (Jackson & Westlind-Danielsson, 1994). More specifically, D2 receptors can be found at low densities throughout the brain, at medium densities in parts of the hippocampus, but only at high density in the dorsal striatum (Palomero-Gallager et al., 2015).

Additionally, the processing of partially temporally predictable stimuli was compared to that of fully temporally predictable stimuli. More specifically, one stimulus block incorporated inter-stimulus intervals (ISIs) of 400ms and 1200ms (termed short and long trials, respectively). This constituted the partially temporally predictable condition. Another block used a fixed ISI of 600ms (termed medium trials), constituting the fully temporally predictable condition. As previously highlighted, this is relevant because naturalistic stimuli likely contain some degree of temporal structure (i.e., partial temporal predictability).

Existing evidence provides conflicting hypotheses regarding how the brain might process deviance when timing is somewhat predictable. Firstly, at these timescales, it was shown that estimates of timing may regress to the mean of a sequence (McAuley & Miller, 2007). As such, the ISI for short and long trials might be processed as “early” and “late” relative to this mean. In other words, the rapid appearance of a stimulus in short trials might induce automatic allocation of attention as their early appearance is “surprising”. This would be reflected by an augmented P3a response at short ISIs, as the P3a is sensitive to stimulus-driven disruption of attentional engagement (Polich & Criado, 2006).

Secondly, when tones fail to occur after a short ISI, the timing of the subsequent tone may be treated as fully predictable (i.e., if not a short trial, must be a long trial), or with increased expectation of a tone’s occurrence. This concept is in accordance with the hazard function that describes how the passing of time increases the likelihood of a stimulus or event to occur (Nobre & van Ede, 2018). This would build an expectation of a tone’s occurrence, thus reducing a surprise P3a response. Furthermore, as previously described, the P3b is enhanced when deviants are predictable in time (Schmidt-Kassow et al., 2009; Schwartze et al., 2011). As such, the P3b response might be larger for long ISIs compared to short ISIs.

Finally, in the partially-predictable temporal condition, individuals might produce two separate memory traces—one for long trials, and one for short trials (Sokolov, 1963a, 1963b). This would make both conditions temporally predictable. In other words, the brain would form separate models for the two different ISIs and both would become “expected”. In this case, all three trials would be treated as temporally predictable, leading to similar P3a and P3b responses between the ISI conditions.

Regarding the dopamine manipulations, higher levels of striatal dopamine might further augment the P3a deviance response at short ISIs, as the P3a is thought to be dependent on dopamine signalling (Polich & Criado, 2006), and dopamine increases are related to the processing of novel or surprising stimuli (Bunzeck et al., 2012). However, due to discrepancies in previous literature, it is unclear how the P3b may differ between dopamine conditions: In Schizophrenia, excessive dopamine signalling seems to lead to a reduced P3b response (Bramon et al., 2004; Yeon & Polich, 2003; Ford 1999), yet other studies show that the P3b is influenced by noradrenaline rather than dopamine signalling (Nieuwenhuis et al., 2005; Polich, 2007).

In addition to ERP responses, the current study assessed the spontaneous eye blink rate (sEBR) derived from continuous resting-state EEG recordings. Many studies have shown a relationship between sEBR and striatal dopamine levels, (i.e., individuals blink more when striatal dopamine levels are higher) (e.g., Sadibolova, Monaldi, & Terhune, 2022; Jongkees & Colzato, 2016; Kowal, Colzato, & Hommel, 2011; Zhang et al., 2015). Other studies have failed to show a relationship between striatal dopamine levels and sEBR (Sescousse et al., 2018; Dang et al., 2017). Nevertheless, sEBR is readily measurable by utilising frontal EEG electrodes; and to our knowledge there are no other non-invasive techniques available to measure endogenous dopamine levels. Therefore, although there is some debate, sEBR might be useful in observing how pharmacological manipulations influenced striatal dopamine levels as in the current study.

Having an outcome measure of the dopamine manipulations is advantageous considering that if dosages of D2 receptor drugs are too low, they can have the opposite to expected effects on dopamine signalling—i.e., agonists decrease dopamine, whilst antagonists increase dopamine (Frank & O’Reilly, 2006; Herrera-Estrella et al., 2005). This occurs due to low dosages stimulating presynaptic autoreceptors rather than postsynaptic D2 receptors, which have opposing effects on neurons (Frank & O’Reilly, 2006). In an attempt to avoid presynaptic effects in the current study, dosages well above the therapeutic starting dosage for PD and Schizophrenia were administered (Del Dotto & Bonuccelli, 2003; Chung et al., 2012); and aligned with dosages shown to influence cognition in previous human studies (Nandam, Hester & Wagner, 2013; Park et al., 2012).

In summary, the main aim of the present study was to investigate the influence of striatal dopamine manipulations on deviance processing when temporal predictability was systematically altered. Regarding the temporal aspects of the study, the available literature led to three main hypotheses that are not mutually exclusive: (1) Short ISIs “surprise” and therefore disrupt the attention system, leading to an augmented P3a response; (2) Long ISIs are processed as fully temporally predictable, and therefore are preferentially processed (increased P3b amplitude for long compared to short ISIs); (3) Two separate memory traces are formed to deal with the two ISIs, making them both fully temporally predictable – meaning that no differences should be observed between conditions. It is likely that dopamine influences the P3a, especially in short trials. However, it remains unclear how the dopamine manipulations may influence the P3a and P3b between conditions. These hypotheses were investigated using an auditory oddball task to simultaneously measure the P3a and P3b subcomponents of the P300.

## 2 Results

Participants performed the auditory oddball task in two blocks (order counterbalanced between participants): The temporally predictable block was comprised of tones with an ISI of 600ms (medium trials); and the semi-predictable block contained trials with ISIs of 400ms and 1200ms (short and long trials, respectively).

One participant experienced an adverse reaction to *amisulpride* and withdrew before completing the study. Another participant withdrew before completing all three sessions. As a result, the remaining data from these participants were not included in analyses, leaving a total of 29 participants for further analysis.

### 2.1 Eye blink rate

One participant was removed from sEBR analysis due to showing an sEBR more than 1.5 times higher than the interquartile range for the antagonist group. There was a significant linear relationship between dopamine level and sEBR; lowest, medium, and highest sEBR rates were found in the antagonist, placebo, and agonist conditions, respectively; F(1,27) = 7.242, p < .05, ηp^2^ = .211; see Figure 1. There was no significant curvilinear relationship between sEBR and dopamine manipulation (p = .926). This demonstrates that in the present study, *amisulpride* and *cabergoline* decreased and increased striatal dopamine, respectively.

**Fig. 1:**
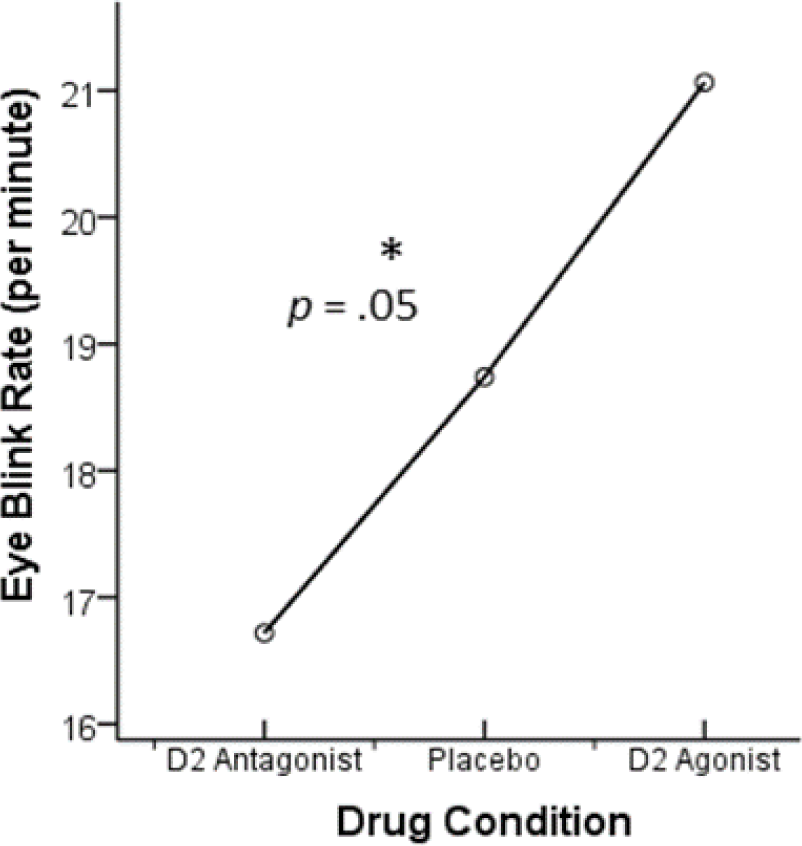
Striatal dopamine reduction using a D2 antagonist (amisulpride) reduced spontaneous eye blink rates, whereas a dopamine agonist (cabergoline) increased sEBRs, relative to placebo. Hence dopamine level is positively related to sEBR using these drugs.

### 2.2 ERPs

Due to experimenter error, EEG triggers were not saved for two participants. The EEG data from three participants were too noisy to distinguish ERPs. As such, these participants were excluded from ERP analyses, leaving a total of 24 participants for further analysis. Via visual inspection of the difference topographies for each condition, it was ascertained that the P3a and P3b had fronto-central and centro-posterior distributions (see supplementary information). As such, the anterior cluster of electrodes consisted of Cz, Fz, Fcz, C1, C2, FC1, FC2, F1, F2 (corresponding to the P3a); and the posterior cluster of CPz, CP2, CP1, Pz, POz, PO3, PO4, P4, P3 (corresponding to the P3b). Visual inspection of the resulting ERPs suggested a time window of 250-320 ms for the P3a response, and 300-500ms for the P3b response, encapsulating the standard and deviant peaks across conditions. For each condition, averaged ERPs consisted of 20-30 clean trials (Cohen & Polich, 1997).

The main effect of ISI length on the P3a response was significant: F(2, 46) = 13.465, *p* < .001, *ηp^2^* = .369. Tests of simple effects demonstrated that P3a responses were significantly larger in the short versus long ISI condition (t(23) = 5.644, *p* < .001 *d* = 1.153), and P3a responses were larger for the medium ISIs than long ISIs (t(23) = 2.897, *p* = .008 *d* = .591). The difference between short and medium ISIs did not reach statistical significance (*p* = .041) following Bonferroni adjustment of the alpha value (*p* < .0167). See Figure 2.

**Fig. 2:**
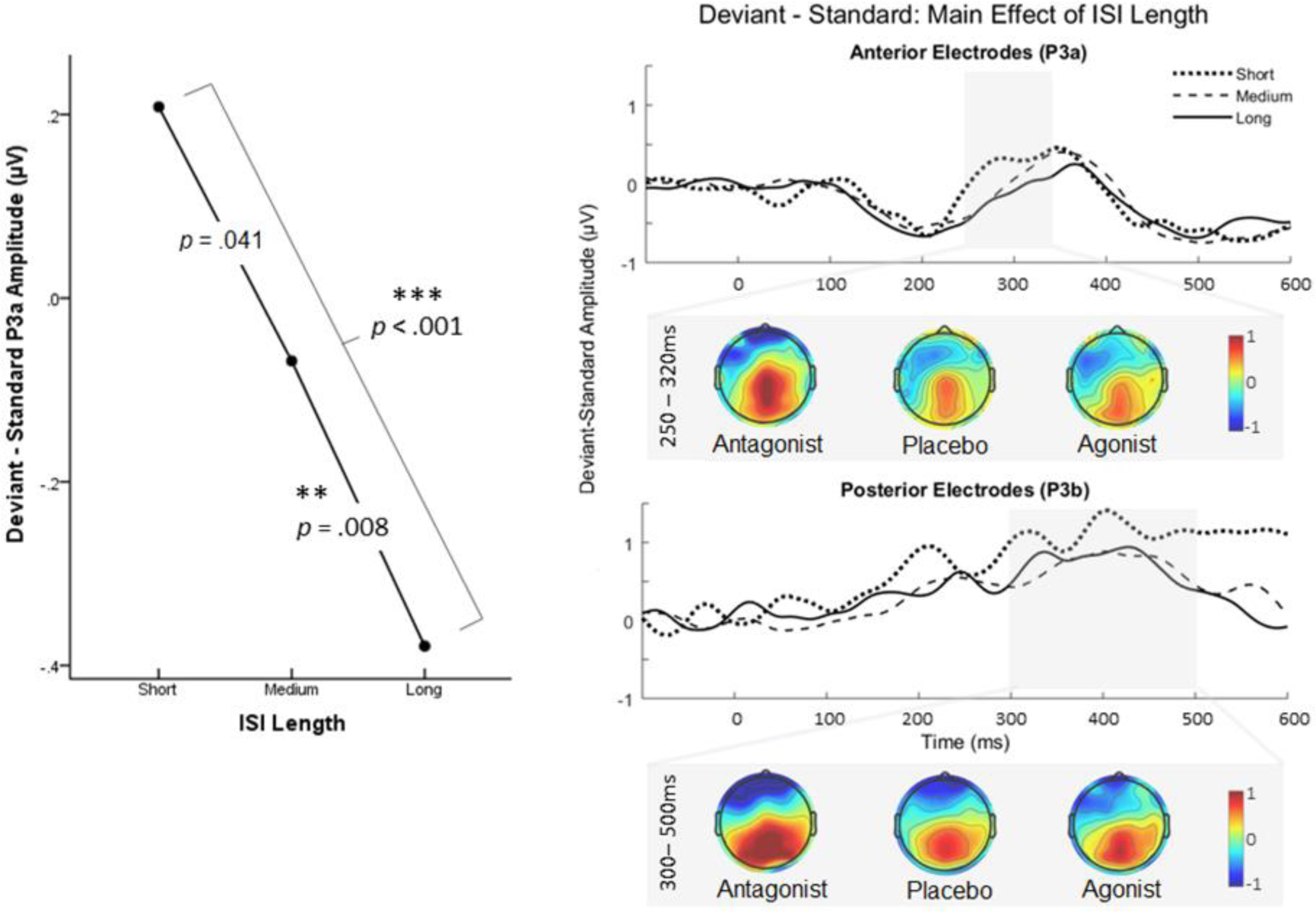
The short ISI was related to a larger P3a effect (left). ERPs and topographies from anterior electrodes and the time window associated with the P3a (top right) and from posterior electrodes and the time window associated with the P3b (bottom right). For graphical representations, an additional low pass filter (25Hz) was applied to ERPs.

The main effect of the drug on the P3a response was also significant: F(2, 46) = 7.933, *p* = .001, *ηp^2^* = .256. Tests of simple effects identified that P3a responses were larger in the placebo condition compared to the agonist (t(23) = 4.088, *p* < .001 *d* = .611) and the antagonist (t(23) = 3.081, *p* = .005 *d* = .632). The Bonferroni-adjusted alpha value was *p* < .0167. See Figure 3.

**Fig. 3:**
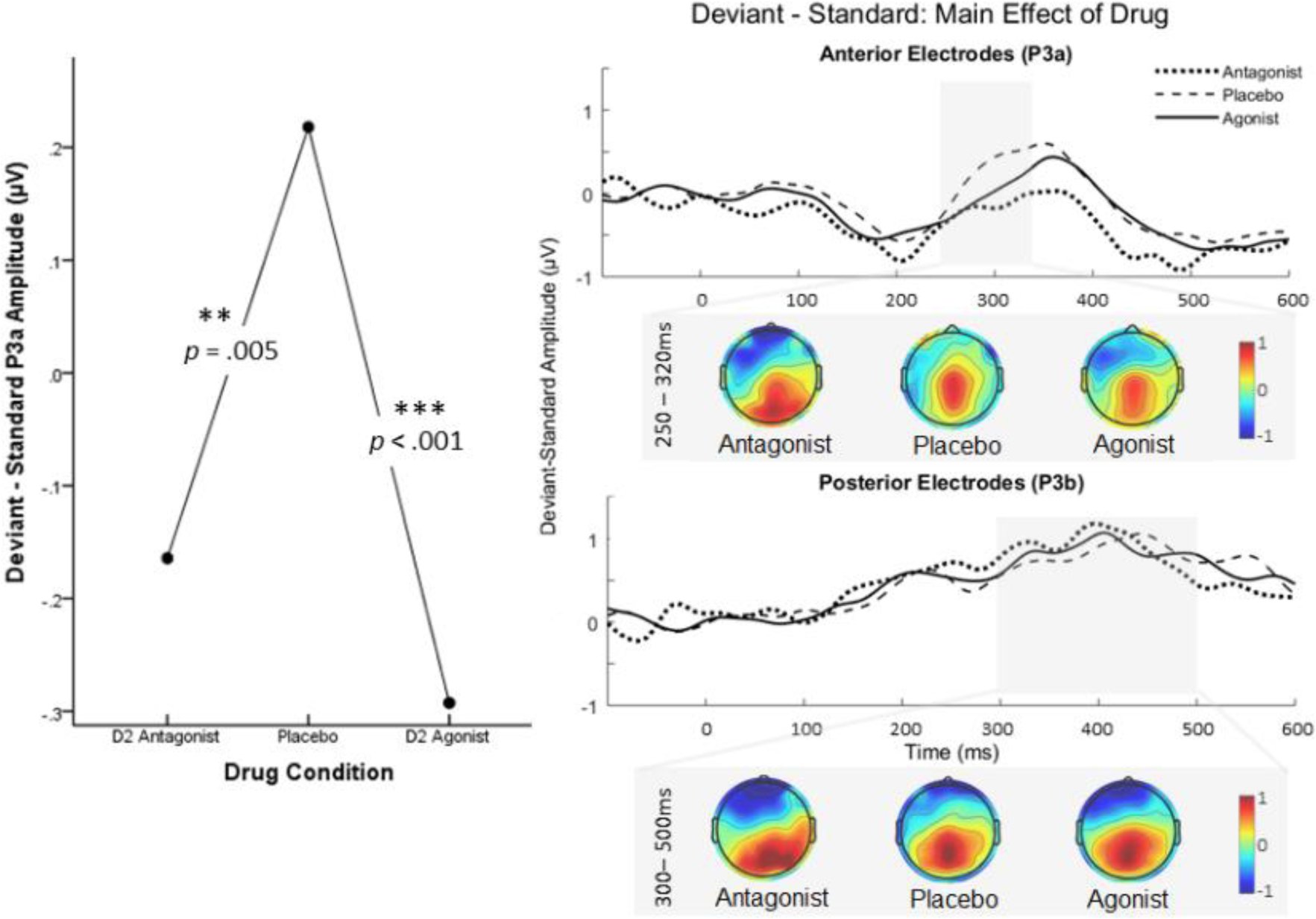
Placebo produced larger P3a effects than both the D2 agonist and antagonist (left). ERPs and topographies from anterior electrodes and the time window associated with the P3a (top right) and from posterior electrodes and the time window associated with the P3b (bottom right). For graphical representations, an additional low pass filter (25Hz) was applied to ERPs.

The interaction between drug and ISI length was significant for the P3a response: F(4, 92) = 3.569, *p* = .002, *ηp^2^*= .163. According to planned tests of simple effects, compared to placebo, P3a responses were smaller at long ISIs under the D2 agonist (t(23) = 3.897, *p* = .001 *d* = .795), but not for the antagonist (*p* = .091). There were no significant differences when comparing the antagonist or agonist with placebo at short ISIs (*p* = .012 and *p* = .118, respectively); or at medium ISIs (*p* = .906 and *p* = .519, respectively). The Bonferroni-adjusted alpha value was *p* < .008. See Figure 4.

**Fig. 4:**
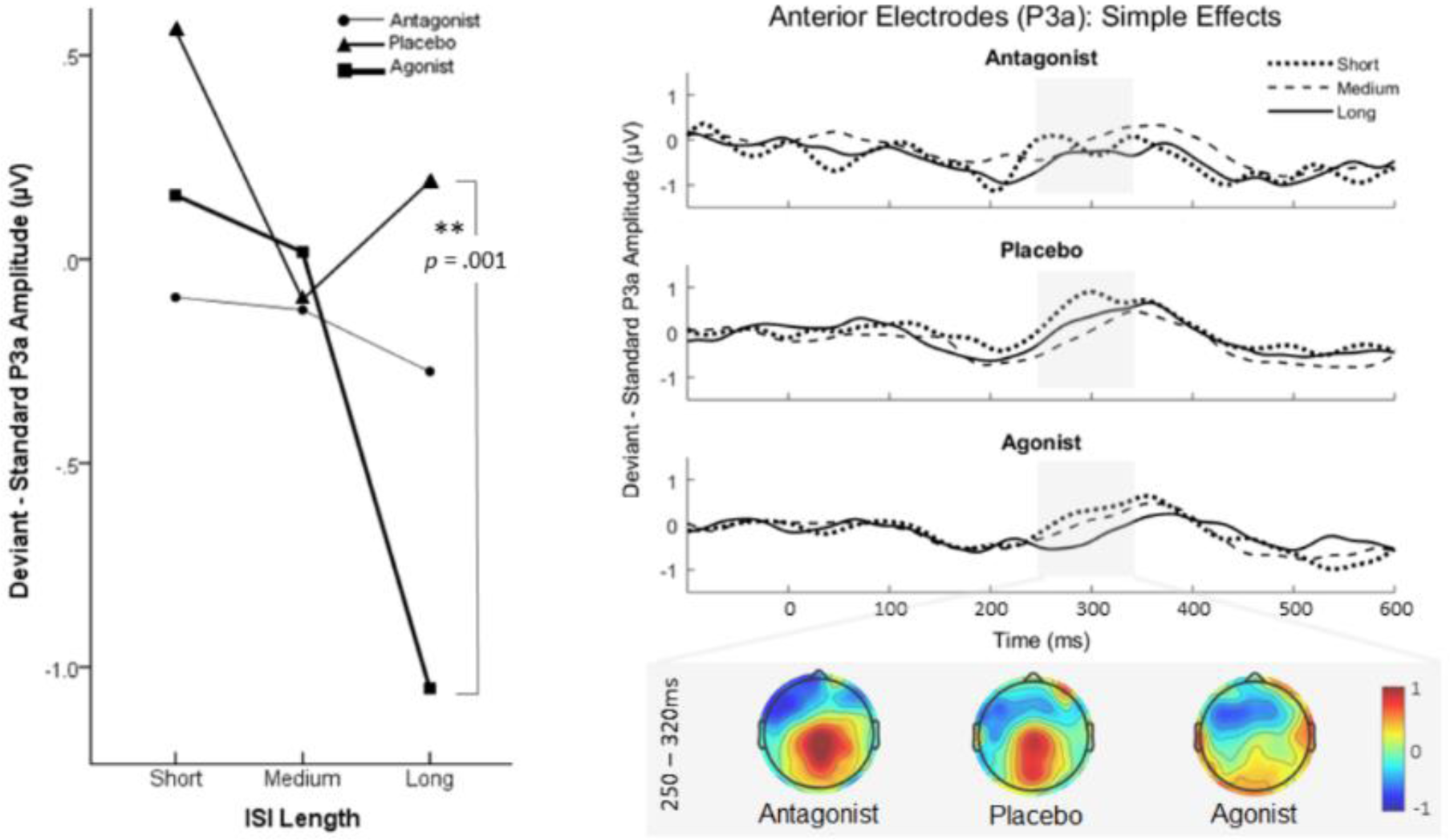
In the agonist condition the P3a effect was larger than in the placebo condition for long ISIs (left). ERPs and topographies from anterior electrodes and the time window associated with the P3a for each drug condition (right). For graphical representations, an additional low pass filter (25Hz) was applied to ERPs.

For the P3b response, neither the main effect of ISI length (F(2, 46) = 2.532, *p* = .091, *ηp^2^* = .009), drug (F(2, 46) = .187, *p* = .830, *ηp^2^*= .008), nor the interaction (F(4, 92) = .775, *p* = .573, *ηp^2^*= .031) were statistically significant.

## 3 Discussion

The present study investigated the influence of striatal dopamine manipulations on deviance processing under different degrees of temporal predictability. The P3a and P3b subcomponents of the P3 ERP complex were taken as indices for the quality of cognitive processes associated with the predictive adaptation to a dynamic auditory environment. Spontaneous sEBR was assessed to verify the effects of pharmacological manipulations on striatal dopamine signalling.

sEBR was highest in the D2 agonist condition, and lowest in the D2 antagonist condition. sEBR rates are thought to be directly linked to striatal dopamine levels (Agostino et al., 2008; Kleven & Koek, 1996; Kowal et al., 2011; Zhang et al., 2015). Therefore, the current results indicate that *cabergoline* and *amisulpride* were likely to have increased and decreased striatal dopamine, respectively (relative to placebo). This finding supports previous studies where persons suffering from schizophrenia exhibited reduced sEBRs when medicated with a D2 antagonist (Adamson, 1994; Karson et al., 1981; Kleinman et al., 1984; Mackert et al., 1988), and extends them to show the same effect in healthy individuals under pharmacological intervention.

When collapsed across dopamine manipulations, longer ISIs were related to smaller P3a responses, and shorter ISIs to larger P3a responses (although the difference between short and medium ISIs did not reach statistical significance using the Bonferroni-corrected alpha value). The P3a response marks the initial disruption of attentional engagement, driven by novel stimuli (Comerchero & Polich, 1999; Polich, 2007). As such, these results indicate that stimuli presented after short ISIs were novel or more “surprising” to the attention system – i.e., processed as arriving “too early”. Conversely, tones after long ISIs lacked this novelty aspect, or were processed as “too late” – reflected by smaller P3a responses. This interpretation is supported by evidence that timing regresses to the sequence mean (McAuley & Miller, 2007), meaning that the short and long ISIs would be considered early and late, respectively by the attention system. Furthermore, there is a large body of evidence indicating that the P3a reflects the capture of attention by novel stimuli (Berti, 2008; Escera & Corral, 2007; Horváth et al., 2008; Muller-Gass et al., 2007; Polich, 2007; Schröger et al., 2015).

The finding of smaller P3a amplitudes for longer ISIs is also in accordance with the concept of the hazard function, which describes how the passing of time increases the likelihood of a stimulus/event occurring (Nobre & van Ede, 2018). As per the hazard function, the longer the ISI in this experiment the more time participants would build the expectation of the occurrence of the tone, thus reducing surprise P3a amplitude.

When collapsed across ISI lengths, P3a responses were diminished under both the D2 agonist and antagonist. This inverted-U relationship between striatal dopamine level and P3a response indicates that dopamine over- and under-signalling are detrimental to the processing of novel, unexpected events. In other words, intermediate levels of striatal dopamine are favourable for the P3a response in the same, healthy individuals. This supports previous research of the P3a responses in PD (Seer et al., 2016; Solís-Vivanco et al., 2015; Tsuchiya et al., 2000), and schizophrenia (Bestelmeyer, 2012; Fisher, Labelle, & Knott, 2010); where striatal dopamine is abnormally low and high, respectively. Reduced P3a amplitudes and increased striatal dopamine synthesis capacity have also been observed in individuals at risk of developing psychosis (Mondragón-Maya et al., 2013; van Hooijdonk et al., 2022). As such, problems with temporal processing observed in these disorders are likely related to abnormal striatal dopamine levels, rather than other aspects of the illnesses or their comorbidities.

Regarding the interaction between dopamine manipulation and temporal predictability, the D2 agonist reduced the P3a response compared to placebo. As previously discussed, excessive striatal dopamine in schizophrenia was related to a reduced P3a response (Bestelmeyer, 2012; Fisher et al., 2010). However, here we find that when timing is partially predictable, relatively long ISIs are particularly susceptible to this effect. This finding may be related to previous studies identifying that persons with schizophrenia perceive time intervals as longer than healthy controls (Rammsayer, 1990; Volz et al., 2001; Wahl & Sieg, 1980), even at a millisecond timescale (Carroll et al., 2008). Excessive striatal dopamine might therefore accentuate the perception of “too late” making the ISI “even later”. In this case, the surprise P3a would be even more diminished, as observed here.

In the present study, the P3b was robust to both manipulations of striatal dopamine, and variations in temporal predictability. The robustness to dopamine manipulations was expected, as the P3b is more sensitive to alterations of the noradrenergic system (Nieuwenhuis et al., 2005; Polich, 2007). However, regarding temporal predictability, previous studies found larger P3b amplitudes when timing was more predictable versus less predictable (Schmidt-Kassow et al., 2009; Schwartze et al., 2011). Here we found no difference in the P3b response between more or less temporally predictable conditions.

One explanation for this lack of P3b differentiation could be that separate memory traces were formed for the short and the long ISI. It has been shown that brains build more than one simultaneous mental representation of timing to efficiently process multifaceted patterns of stimuli (Sokolov, 1963b, 1963a). As previously hypothesised, if two representations were built to process the short and long ISIs, both would be treated as equally predictable. In this case, each of the medium, short, and long ISIs would have their own mental model, meaning that the P3b responses would be similar between them (as seen here). However, interpreting null findings must always be approached with caution. Further investigation is necessary to ascertain the parameters under which multiple memory traces are formed.

In this study, we manipulated temporal predictability in a novel manner, by presenting tones at three different ISI lengths. This reflects naturalistic stimuli, as timing is sometimes partially predictable, and can arrive “too early” or “too late”. Conversely, previous studies presented tones at a range of ISIs randomly sampled from a distribution (Schmidt-Kassow et al., 2009; Schwartze et al., 2011). This difference between paradigms is subtle, yet produced a different pattern of electrophysiological effects, thus highlighting the heterogeneity of deviance processing mechanisms, particularly under different degrees of temporal predictability.

Studying the heterogeneity of how temporal predictions are formed – especially the role of dopamine – is important considering diseases such as PD and schizophrenia. Patients with PD and schizophrenia exhibit abnormal temporal processing of both ‘on’ and ‘off’ medication (Carroll et al., 2008; Davalos et al., 2003; Jones et al., 2011; O’Boyle, Freeman, & Cody, 1996). This is likely due to a lack of specificity of dopaminergic treatments, mitigating some cognitive functions whilst worsening others (Jones & Jahanshahi, 2014; Weintraub et al., 2010). Indeed, it is not clear which neurological issues in PD and schizophrenia are caused by the progression of the disease, or by exposure to pharmacological treatments that overstimulate certain receptors or regions of the brain (i.e., due to lack of specificity) (Cools et al., 2001, 2003; Goff et al., 2017; Ho et al., 2011). As such, understanding the interaction between specific dopamine receptors in specific cognitive functions is important for the improvement of treatment specificity.

## 4 Conclusion

In conclusion, in the present study, temporal predictability was manipulated in a manner which reflects naturalistic stimuli (i.e., partially predictable). Results demonstrate that in healthy individuals, increased dopamine signalling in the striatum via a D2-receptor agonist can interfere with temporal processing. This could help to explain why in dopaminergic disorders such as PD, treatments that target D2 receptors might worsen certain cognitive symptoms (Cools et al., 2001, 2003). Cognitive symptoms are highly debilitating, and in some patients negatively affect quality of life more than physical symptoms (Caballol, Martí, & Tolosa, 2007). As such, these findings point towards further research into the specific roles of dopamine subsystems in temporal processing, with the wider view of improving the treatment of dopaminergic disorders.

## 5 Methods and materials

### 5.1 Design

A double-blind, placebo controlled, repeated measures, triple crossover design was employed. Participants attended the lab three times each separated by at least one week. At each visit, participants orally self-administered either *cabergoline* (1.25mg), *amisulpride* (400mg), or placebo (sugar pill).

### 5.2 Participants

Thirty-one participants were deemed eligible to participate in the full study (2 left-handed; 16 female; mean age 22.4 years; standard deviation 3.2 years). Details of the medical health screening are included in supplementary information. All participants gave written informed consent, and received financial compensation for their participation. The study was approved by the University of Manchester Research Ethics Committee (ref. 14194), and was further approved by the UK Health Research Authority due to the use of a National Health Service site (Salford Royal Hospital).

### 5.3 Procedure

#### 5.3.1 Pre-Task

Both of the active drugs reach peak plasma levels around two hours following oral administration (Del Dotto & Bonuccelli, 2003; Mauri et al., 2014). Following drug administration, participants remained in a neutral environment for two hours before testing procedures began. During this time, EEG scalp electrodes were connected.

EEG was recorded using sixty-four *Easycap* scalp electrodes (easycap.de), at a sampling rate of 1000Hz, and amplified by a *Brainvision BrainAmp DC plus MR* amplifier. Spontaneous eye blinks were then measured for nine minutes. During this time, participants were asked to rest (without closing their eyes) whilst the EEG electrodes recorded their “resting activity”. The auditory oddball paradigm was administered 2.5 hours following drug administration following a 30-minute visual attention task as part of a separate investigation.

#### 5.3.2 Oddball Task

Auditory stimuli in the form of a sequence of tones were presented via headphones whilst the participant was seated in a lit room with eyes open. Continuous EEG was recorded throughout. Stimuli were 600Hz and 660Hz pure tones with a duration of 300ms. One of the tones was presented with a frequency of 75% (the standard) and the other with a frequency of 25% (the deviant). Which of the tones was the standard versus the deviant was counterbalanced between participants. In one block, tones were presented with a 600ms ISI (predictable condition), whilst in another block, ISIs were randomly 400 and 1200ms (semi-predictable condition). Block order was counterbalanced between participants. See Figure 5 for a schematic diagram depicting the experimental paradigm. Blocks consisted of 520 tones, making each around 8 minutes in duration. Between blocks, participants were provided with a break of around 5 minutes. In order to promote attention to the stimuli, participants were instructed to have their eyes open and to count the number of occurrences of the deviant tone and report the value to the experimenter at the end of the block.

**Fig. 5:**
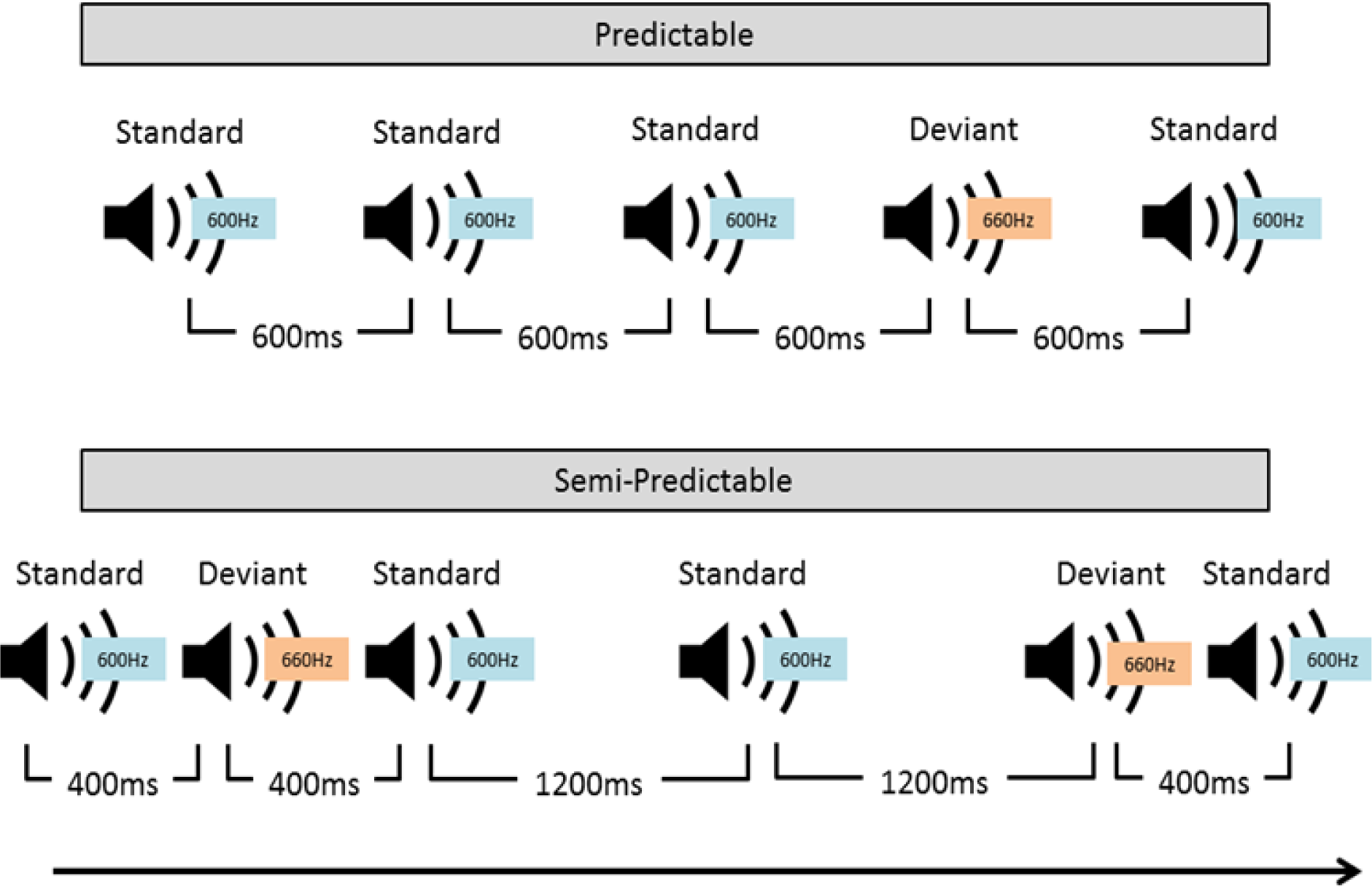
Schematic diagram of example trials from the auditory oddball paradigm. In this example, the first block presents tones at a fully predictable ISI of 600ms (upper panel), with the second block containing ISIs of either 400 or 1200ms (semi-predictable, lower panel).

### 5.4 Analyses

#### 5.4.1 Blink rate

A value of blinks-per-minute was calculated for each participant and visit. Using frontal electrodes, continuous EEG data were downsampled to 200Hz and filtered offline. Independent component analysis (ICA) was performed using the jader function from the EEGLAB open-source toolbox (Delorme & Makeig, 2004). The component corresponding to eye blinks was identified via visual inspection. Blinks were quantified using a custom MATLAB script (The Mathworks, Inc.) to count the number of prominent voltage deflections in the identified component. Contrast analysis was employed to test for the predicted linear relationship between dopamine manipulation and sEBR.

#### 5.4.2 ERPs

Before ERP extractions, EEG recordings were pre-processed offline using SPM12 software (Statistical Parametric Mapping, UCL, England); electrodes were re-referenced to a whole-scalp reference, low-pass filtered at 40Hz, downsampled to 200Hz, and then high-pass filtered at 0.1Hz. ERP epochs were then defined as −100ms to 600ms relative to the onset of the target display, were averaged, and baseline-corrected relative to the 100ms pre-stimulus time window. Trials contaminated by artefacts (including eye-blinks/movements) were excluded from analyses, detected as events recorded at any of the electrode channels exceeding 75μV relative to the pre-stimulus baseline. P300 amplitude has been found to stabilise at around 20 trials per condition (Cohen & Polich, 1997). As such, averages were only considered for analysis if they contained at least 20 clean trials (i.e. without artefacts).

Deviant minus standard difference waves were created separately for each drug condition (antagonist, placebo, and agonist) and ISI length (short, medium, and long). Difference waves were extracted from two separate electrode clusters according to the P300 literature: Anterior (corresponding to the P3a) and posterior (corresponding to the P3b) (Polich, 2003, 2007). The electrodes included in clusters were defined by visual inspection of deviant minus standard scalp topographies. Time windows for these inspections were based on broad time windows for the P3a and P3b (250-350ms and 300-500ms, respectively) as guided by the extensive P300 literature (Comerchero & Polich, 1999; Donchin, 1981; Polich, 2003; Polich & Criado, 2006). Using the resulting ERPs, more specific time windows for the P3a and P3b were identified via visual inspection, defining intervals which encompass the effects observed across the conditions. At these time windows, amplitudes of the difference waves were averaged, providing single values for the “P3a response” and “P3b response” for each condition.

To test for differences between conditions, repeated-measures ANOVAs were performed separately for the P3a and the P3b responses. The ANOVAs consisted of two variables, each with three levels: Drug (antagonist, placebo, and agonist), and ISI length (short, medium, and long). Significant main effects and interactions were further investigated using within-subjects t-tests (tests of simple effects) with Bonferroni-adjusted alpha values. In order to investigate how the dopamine manipulation affected temporal processing, t-tests compared each drug to placebo separately for each ISI condition (tests of simple effects for the interaction).

## 9 Acknowledgements

The authors would like to thank the participants for their time and commitment to the study, Tim Rainey for technical support, and Dr. Christopher Kobylecki for performing the health screenings and providing medical support. The research was funded by the Biotechnology and Biological Sciences Research Council Doctoral Training Partnership (BBSRC-DTP).

G.A.W. was supported in part by the Centro Basal FB0008.

## 7 Supplementary information

### 7.1 ERP Extractions

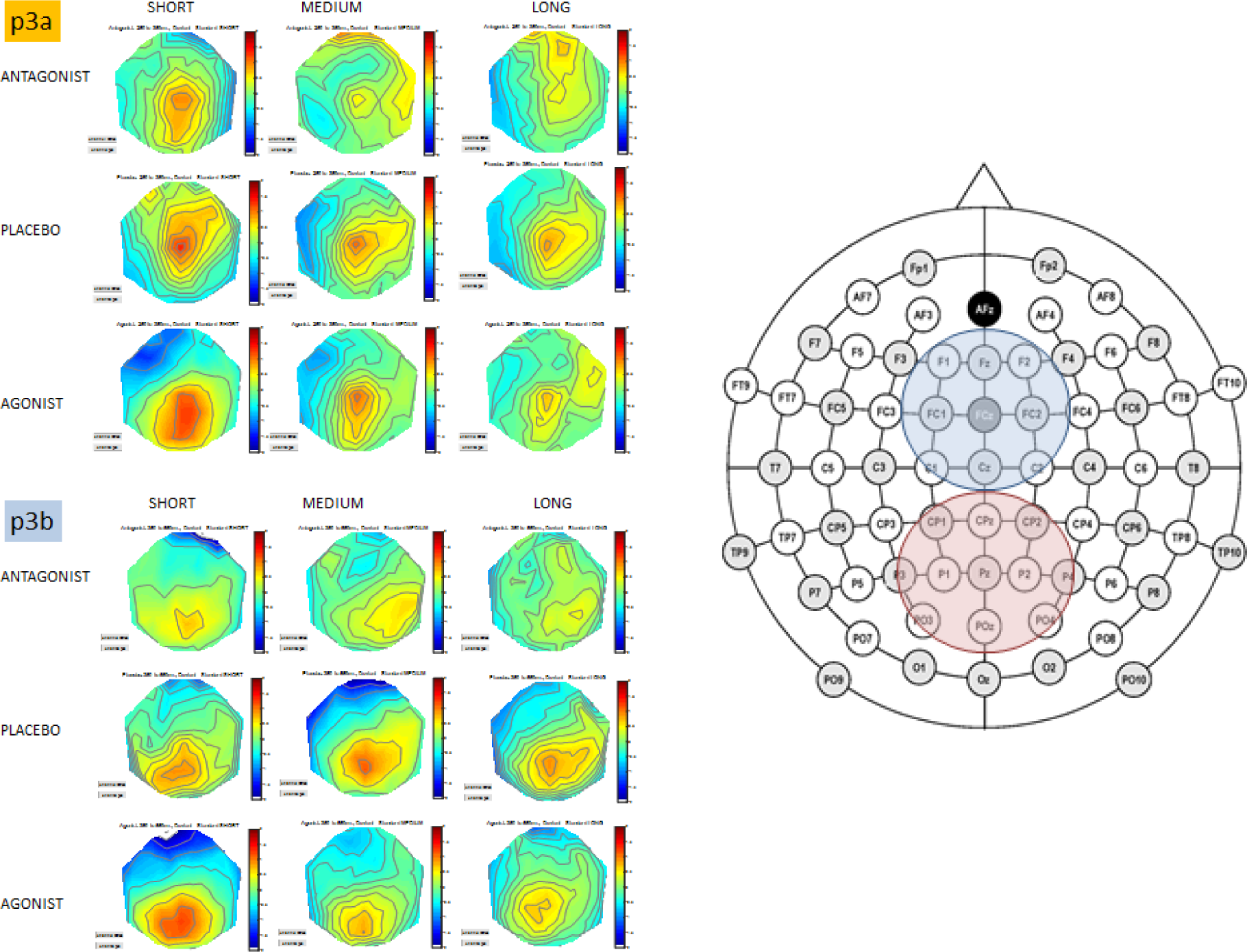
Scalp topographies for typical P3a and P3b time windows (250-300ms and 300-500ms, respectively). This identified two electrode clusters along the midline, each consisting of an equal number of electrodes (anterior P3a in blue, posterior P3b in red).

### 7.2 Health Screening

Thirty-one individuals attended further health-screening by a clinician. All were deemed fully eligible to participate in the study. It was required that participants were non-smokers with normal or corrected-to-normal vision, were not colour blind, and were fluent in the English language. It was also required that they had none of the following: History of significant head injuries or seizures, diagnosis of any neurological or psychiatric condition, history of drug or alcohol abuse, use of psychotropic medication within the past six months, use of dopaminergic drug within the past month (or lifetime use exceeding three months), or history of heart problems. It was required that females were not pregnant, breast feeding, or trying to conceive.

As amisulpride can cause small changes in heart function (prolongation of the QTc interval) (Täubel et al., 2017), a clinician ensured that heart rate, blood pressure, and electrocardiogram measures were within a healthy range. Clinicians also took a brief medical history focusing on cardiac abnormalities. Participants’ general practitioners were informed of the study, and were asked to notify the research team of any potential concerns.

### 7.3 Precautions and Aftercare

To avoid possible drug interactions, participants were asked to refrain from taking prescription or non-prescription medications for the duration of the study, with the exception of the contraceptive pill and paracetamol (unless discussed with a study clinician). They were also asked to refrain from consuming alcohol 24 hours prior to participation days, and to consume their typical amounts of caffeine on participation days.

At the end of each visit, participants received a ‘contact card’, stating that the carrier had participated in a research study involving dopamine agonists and antagonists. The card also included contact phone numbers for members of the research team, as well as the participant’s unique identification number for emergency unblinding. Participants were instructed to call one of these numbers if they experienced any unusual sensations after their visit.

